# A method for differential expression analysis and pseudo-temporal locating and ordering of genes in single-cell transcriptomic data

**DOI:** 10.1101/2022.12.21.521359

**Authors:** Bao Zhang, Hongbo Zhang

## Abstract

Identification of differentially expressed genes (DEGs) is a pivotal step in single-cell RNA sequencing (scRNA-seq) data analysis. The sparsity and multi-model distribution of scRNA-seq data decides that the traditional tools designed for bulk RNA-seq have several limitations when applied to single-cell data. On the other hand, tools specifically for DEGs analysis of scRNA-seq data normally does not consider the high dimensionality of the data. To this end, we present DEAPLOG, a method for differential expression analysis and pseudo-temporal locating and ordering of genes in single-cell transcriptomic data. We show that DEAPLOG has higher accurate and efficient in DEGs identification when compared with existing methods in both artificial and real datasets. Additionally, DEAPLOG can infer pseudo-time and embedding coordinates of genes, therefore is useful in identifying regulators in trajectory of cell fate decision.

## Introduction

Recently developed single-cell RNA sequencing (scRNA-seq) technologies have enabled high-throughput whole-transcriptomic profiling at single-cell resolution, and hundreds of bioinformatics tools have been developed to process, analyze, and interpret the results from scRNA-seq data, such as the toolkits Scanpy and Seurat, making it possible to dissect heterogeneities within cell states, developmental stages, and disease status^1,2^. One key step of analyzing scRNA-seq data is to identify differentially expressed genes (DEGs). In contrast to bulk RNA-seq, scRNA-seq data has two special characteristics. One is the sparsity, which is indicated by “dropout events” (i.e. a large proportion of zeros in single-cell expression matrix)^3^. The other is that the expression distribution of gene is multi-modal^4^. Both cause challenges in identifying DEGs.

Although most common methods developed for bulk RNA-seq have been used to identify DEGs in scRNA-seq data, the drawback of these methods is that traditional probability distributions do not consider the two special characteristics of scRNA-seq data^5^. Driven by this, a few methods have been designed specifically for scRNA-seq data. For example, MAST, DEsingle, DECENT and SCDE employed mixture model to quantify “dropout events” but not multimodal distribution^6-9^; whereas, D3E, monocle 2, NBID and scDD consider the possibility of multi-modal distribution of scRNA-seq data but not sparsity^10-13^. The only tool ZIAQ considers both factors^14^, yet, as most of other methods do, is limited to two-class comparisons and cannot adapt for multiple conditions, which is often the case in complex cell status and experimental designs. Additionally, recent comparisons found that the current scRNA-seq-specific DEG identification approaches do not necessarily perform better than those for bulk RNA-seq^15^.

One possible explanation for this is that all of these methods are based on probabilistic models and ignore the advantages of high throughput of cells in scRNA-seq. In contrast to the small sample size of bulk RNA-seq data, scRNA-seq data have thousands of cells to characterize the expression patterns of each gene, making it possible to identify the cell signature and perform cell signature enrichment analysis (CSEA) for any given gene.

Driven by these factors, we propose a new method named DEAPLOG (Differential Expression Analysis and Pseudo-temporal Locating and Ordering of Genes) for identifying DEGs in scRNA-seq data. Comparisons with existing methods show that DEAPLOG has higher sensitivity, specificity, and efficiency on both artificial and real datasets. DEAPLOG is flexible enough to be extended to complicated conditions, such as multiple cell-clusters and experimental conditions. DEAPLOG can also be used to infer pseudo-time and embedding coordinates of genes, which is useful in identify key regulators along the trajectory of cell fate decision.

## Results

### Framework of DEAPLOG

The DEAPLOG has two functions for analyzing single-cell sequencing data: one is to identify DEGs for cell clusters or other experimental conditions, and the other is to infer the pseudo-time and embedding coordinates of genes (Fig. 1). For identifying DEGs for cell clusters, DEAPLOG first identifies cell signatures for each gene, and then performs CSEA on cell clusters (see methods). After that, we obtain the cell cluster to which the cell signature of each gene belongs to, and in turn, identify indirectly DEGs of each cell cluster based on the enriched significance score. The cell clusters can be originated from different cell types but also other experimental conditions, such as developmental stages and pathological states. For pseudo-temporal locating and ordering of genes, DEAPLOG infers the pseudo-time and embedding coordinate of a gene based on the pseudo-times and embedding coordinate of its cell signature obtained at the previous step (refer to “Methods” section for details). In conclusion, DEAPLOG is a tool not only for identifying DEGs for cell clusters, but also for inferring the pseudo-time and locating embedding coordinate of genes, which is great helpful for studying cell inherent heterogeneity and dynamic changes, as well as exploring potential key players that drive the status changes, such as transcription factors (TFs) that involve in cell lineage differentiation during embryonic development.

**Fig. 1.**
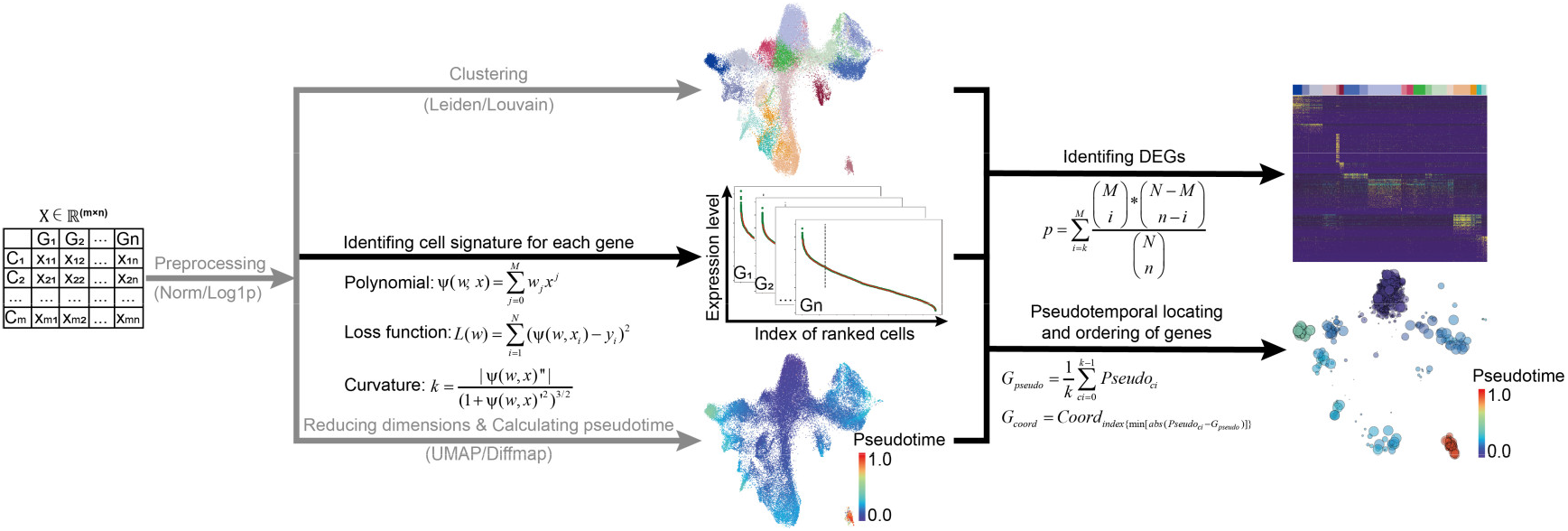
the workflow of DEAPLOG. The schematic of the workflow of DEAPLOG for DEGs identification and pseudo-temporal locating and ordering of genes. A scRNA-seq dataset of human hematopoietic lineages were used for method visualization. The processes indicated by black solid lines and arrows are executed by DEAPLOG, while the processes indicated by gray solid lines and arrows are executed by other software and used as input data for DEAPLOG.

### DEGs identification by DEAPLOG

To further elucidate the principle of DEAPLOG in DEGs identification, we use a human scRNA dataset from hematopoietic stem/progenitors (HSPC) and its progeny cells as an example considering that the HSPC lineage has well-studied known cell composition^16^. Leiden clustering shows that this data contains 21 cell clusters from three main lineages (lymphoid, myeloid and erythroid) derived from HSPC, which are all able to be distinguished with classic markers (Supplementary Fig. 1A, B). For example, B cells can be well-identified using the CD27 marker^17^ (Fig. 2a). To validate DEAPLOG, we first identify the cell signature for *CD27*. The results show that the top 426 cells identified as cell signature for *CD27* are almost all belong to the cluster of B cells, which proves the validity of cell signature analysis (Fig. 2b). Then, we perform CSEA for *CD27* and found that it is a very significant DEG (adjusted p-value = 2.2e-299 and ratio = 0.937) which is only enriched in B cells (Fig. 2c). In addition to CD27, VPREB1 also plays an important role on B cell differentiation^18^. We first find that *VPREB1* is expressed in B cells, together with its precursor cycling B cells and early lymphoid cells (EarlyLymp) (Fig. 2d). We then perform DEGs analysis using DEAPLOG. The top 4,195 cells are identified as cell signature for *VPREB1* and indeed almost all these cells belong to the three cell clusters: B cells, cycling B cells and EarlyLymp (Fig. 2e). The result of CSEA by DEAPLOG also shows that VPREB1 is significantly enriched in B cells (adjusted p-value = 8e-300 and ratio = 0.446), EarlyLymp (adjusted p-value = 8e-300 and ratio = 0.304) and cycling B cells (adjusted p-value =8e-300 and ratio = 0.205) (Fig. 2f). This confirms that DEAPLOG can be utilized to identify DEGs shared by multiple clusters. As a negative control, *GAPDH*, a housekeeping gene, is expressed in almost all cells and the results of cell signature and CSEA by DEAPLOG also prove that *GAPDH* is a wildly-expressed gene in HSC lineages (Fig. 2g-i).

**Fig. 2.**
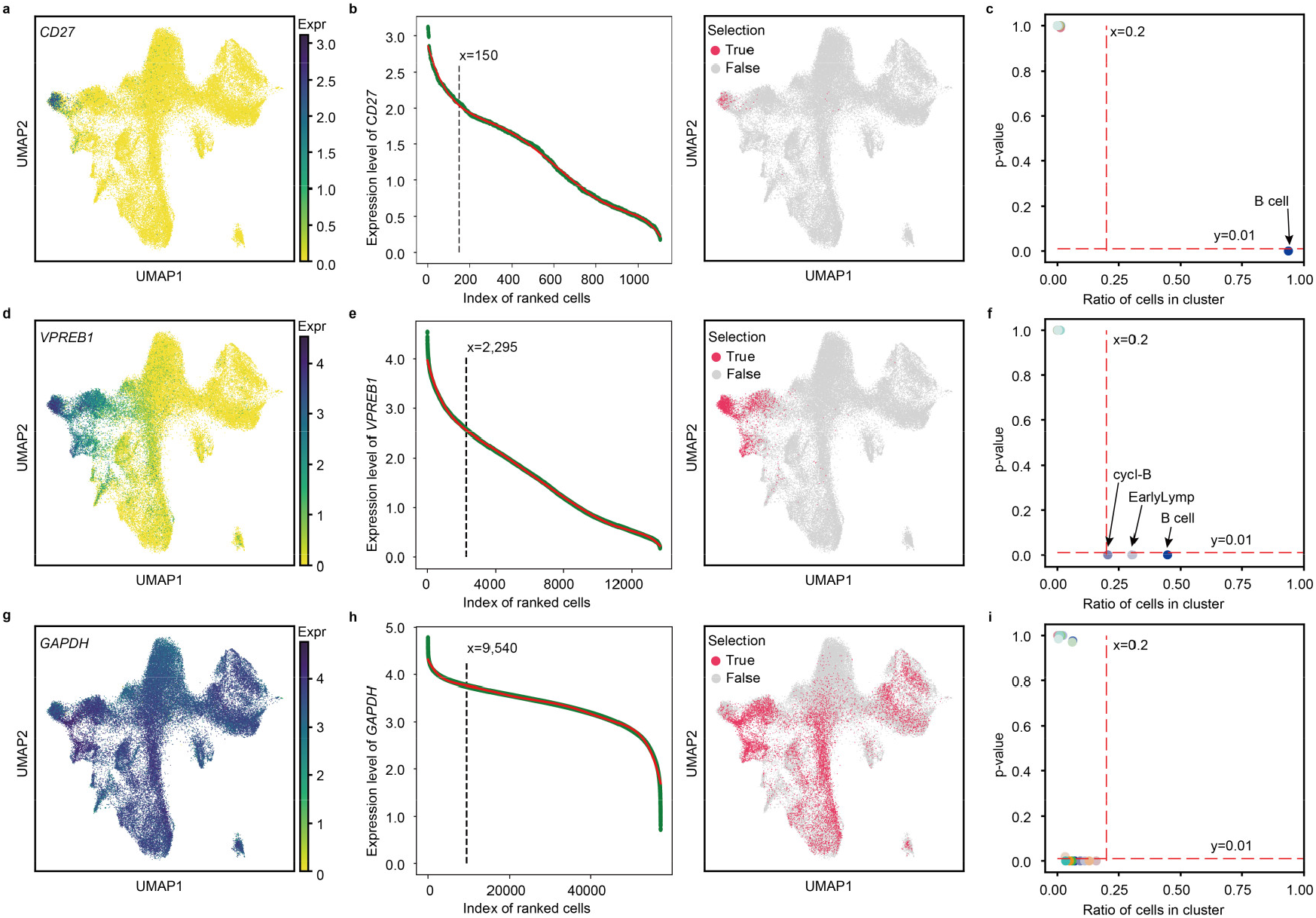
Illustration of DEGs identification by DEAPLOG. Illustration of the performance of DEAPLOG for DEGs identification by using known marker genes of B cells and the human hematopoietic development data. **a, d, g** UMAP plots showing the normalized and logarithmic expression level of *CD27, VPREB1* and *GAPDH*, respectively. **b, e, h** Scatter (left panel) and UMAP (right panel) plots showing the ranked cells which identified as the cell signatures of *CD27, VPREB1* and *GAPDH*, respectively. Cells located on the left side of the dashed line (left panel) or colored in red (right panel) were considered as the set of cell signatures. **c, f, i** Scatter plot showing the p-value and ratio of *CD27, VPREB1* and *GAPDH* in each cell clusters. Ratio represents the percentage of the cell signatures set for a gene in a cell cluster.

It’s worth noting that if one gene is expressed in more than three cell groups, it is generally considered not a good DEG. For DEAPLOG, we recommend setting the threshold of ratio to 0.2 and the threshold of adjusted p-values to 1e-30 as this can greatly reduce the false positive rate.

### Pseudo-temporal locating and ordering of genes by DEAPLOG

The other function of DEAPLOG is to infer the pseudo-time and embedding coordinates of genes in cell evolution trajectories. To demonstrate the principle of this function, we still use cells from the same data set because the development of HSPC is an excellent model and well-used to explore the regulators driving its lineages differentiation. We focus on B cell developmental lineage, which is composed of four cell clusters: HSC/multipotent progenitor (HSC/MPP), lymphoid MPP (Lymp-MPP), EarlyLymp and B cells (Fig. 3a). *GATA3* is a TF mainly expresses in HSC/MPP (Fig. 3b). We identify the cell signature for *GATA3* by DEAPLOG and find that indeed almost all these cells belong to the HSC/MPP (Fig. 3c). Next, we infer the pseudo-time and embedding coordinate of *GATA3* by DEAPLOG based on the pseudo-time and embedding coordinates of these enriched cells. The result shows that the pseudo-time of *GATA3* is 0.031 and the coordinate of *GATA3* is located at the center of HSC/MPP, suggesting that *GATA3* is an early-stage TF that expressed in HSC/MPP (Fig. 3d). This is consistent with previous studies, which reported that GATA3 regulates hematopoietic stem cell maintenance and cell-cycle entry^19^. Additionally, we calculated the pseudo-time and embedding coordinate of other three TFs: ZNF683, GAS7 and PAX5. Our results show that the pseudo-times of *ZNF683, GAS7* and *PAX5* are 0.085, 0.236 and 0.561, and the embedding coordinates of these three TFs are located at the center of Lymp-MPP, EarlyLymp and B cells, respectively. All consistent with their expected expression patterns (Fig.3e, f). Overall, these results indicate that DEAPLOG can accurately infer the pseudo-time and locate of genes for B cell development, and the pseudo-times and location of *GATA3, ZNF683, GAS7* and *PAX5* suggest that these four TFs are activated at different time points and cell clusters to regulate B cells development (Fig. 3g).

**Fig. 3.**
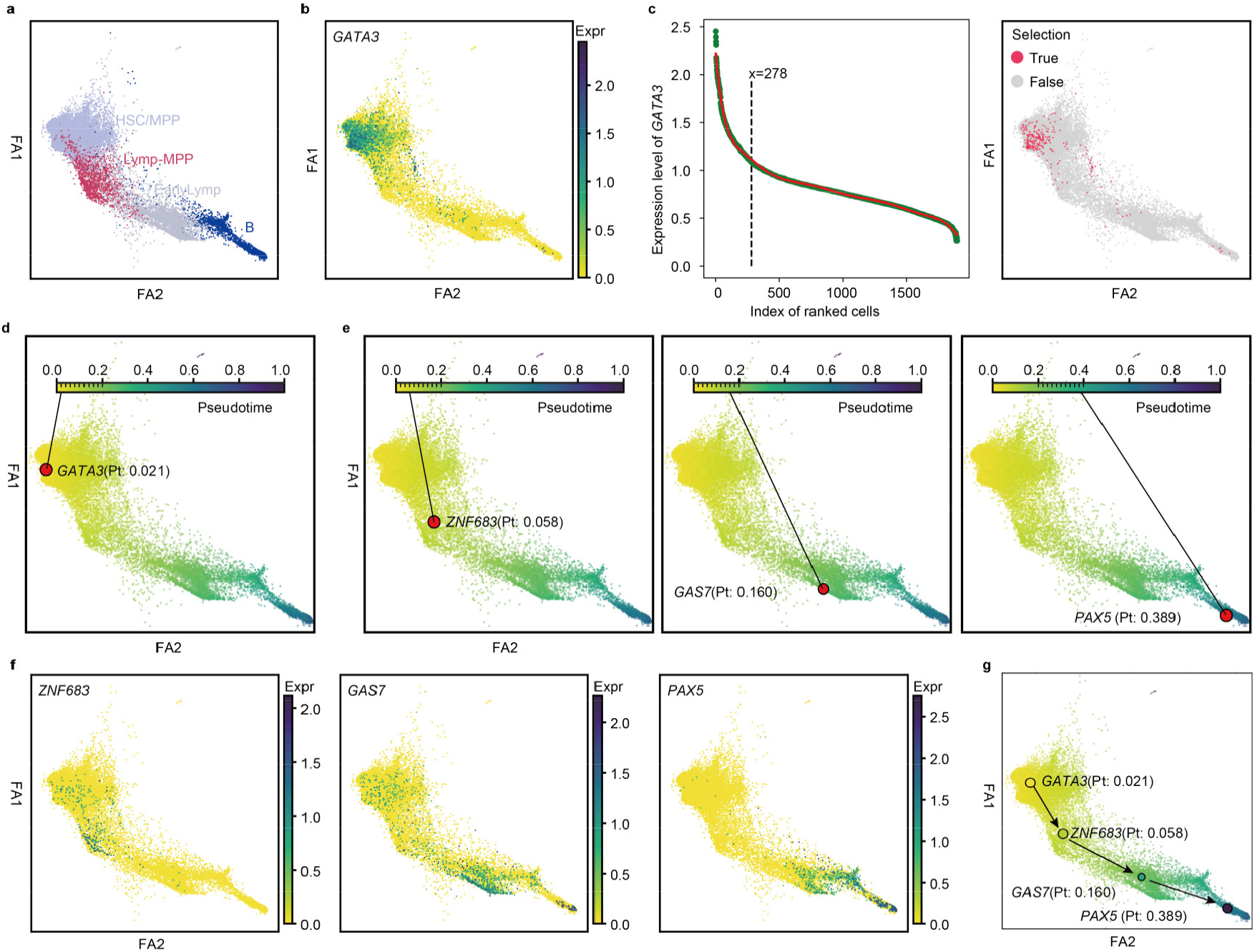
Illustration of pseudo-temporal locating and ordering of genes by DEAPLOG. Illustration of the performance of DEAPLOG for pseudo-temporal locating and ordering of genes by using B cells lineage of the human hematopoietic lineage development. **a** Force-directed graph showing the B development lineage and its cell clusters. **b** Force-directed graph showing the normalized and logarithmic expression level of *GATA3*. **c** Scatter plot (left panel) and force-directed graph (right panel) showing the ranked cells which identified as the cell signatures of *GATA3*. Cells located on the left side of the dashed line (left panel) or colored in red (right panel) were considered as the set of cell signatures, respectively. **d, e** Force-directed graph showing the pseudo-temporal locating and ordering of *GATA3, ZNF683, GAS7* and *PAX5*, separately. **f** Force-directed graph showing the normalized and logarithmic expression level of *ZNF683, GAS7* and *PAX5*, separately. **g** Merged force-directed graph showing the pseudo-temporal locating and ordering of *GATA3, ZNF683, GAS7* and *PAX5*. Solid black arrows represent the direction of B cell development and the Pt value is the pseudo-time of genes.

### Comparison of DEAPLOG with other DEGs identification tools

To evaluate the accuracy of our approach, we first simulate 27 artificial datasets with varying cell number, cell clusters and batches using Splatter in which true DEGs are known (see“Methods”)^20^. Then We apply DEAPLOG and 9 existing methods (Bimod, Logistic regression (LR), MAST, Monocle, Negative binominal (NegBinom), Poisson, SinglecellHaystack (SCHS), t-test and Wilcoxon rank− sum test (Wilcoxon)) on the artificial datasets and compared their accuracy and runtimes^2,6,11,21^. The accuracy of each method was estimated using the area under the receiver operating characteristic (ROC) curve (AUC).

The first 6 datasets are simulated with varying cell numbers from 50 to 5,000 but fixed 2 clusters. In this condition, all methods show comparable good performance except the SCHS, which does not have an AUC value above 0.8 until the total cell number reach 2,500. It’s AUC value even decreased to 0.5 when the total cell number lower than 100. In contrast, the AUC value of DEAPLOG keeps higher than 0.7 even when the total cell number drop to 50 (Fig. 4a and Supplementary Fig. 2). We then compare the performance under different complexity of clusters by simulating 10 datasets of varying cell clusters (2, 5, 10, 25 and 50) with fixed number of cells at either 200 per cluster or 10k in total. To test the effect of data distribution, we also simulated extra 5 datasets of which the cell number in cell clusters follows Poisson, Beta, Gamma, Normal and Uniform distribution, respectively with fixed total 10k cells and 10 clusters in each dataset. Under these conditions, DEAPLOG and SCHS show comparatively high accuracy and stability, while the other methods not only show significantly low accuracy, but also low stability (Fig. 4a and See Supplementary Fig. 3). Worth mentioning that DEAPLOG has higher accuracy than SCHS for the datasets with fixed 200 cells per cluster and at low cluster numbers, again implying that the performance of SCHS is heavily dependent on the total number of cells (Fig. 4a and Supplementary Fig. 3A). Another important factor affecting DEGs identification is batch effect. Therefore, we also applied these methods on 6 artificial datasets of varying batches (1, 2, 4, 6, 8 and 10). Although all methods show high robustness against varying batches, DEAPLOG and SCHS show much higher accuracy than all others (Fig. 4a and Supplementary Fig. 4).

**Fig. 4.**
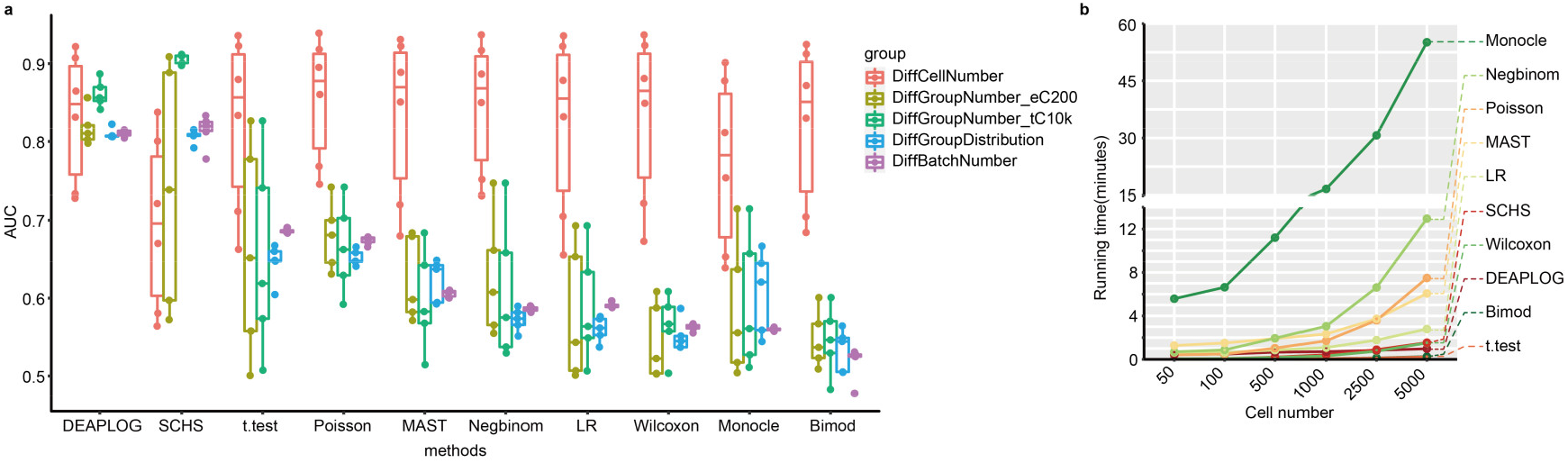
Comparison between DEAPLOG and other DEG prediction methods. A comparison of performance between DEAPLOG and other methods applied on the simulated 27 artificial datasets. **a** boxplot showing the AUC values of different methods applied on different datasets. The datasets belonging to the group of “DiffCellNumber” contains 6 datasets of 50, 100, 500, 1000, 2500 and 5000 cells, respectively. The datasets belonging to the group of “DiffGroupNumber_eC200” contains 5 datasets of 2, 5, 10, 25 and 50 cell clusters, respectively. Each cluster is fixed with 200 cells. The datasets belonging to the group of “DiffGroupNumber_tC10k” contains 5 datasets of 2, 5, 10, 25 and 50 cell clusters, respectively. Each dataset contains 10,000 cells in total. The datasets belonging to the group of “DiffGroupDistribution” contains 5 datasets of 2, 5, 10, 25 and 50 cell clusters with different distribution (Poisson, Beta, Gamma, Normal and Uniform distribution) of cell number per cluster in each dataset. SCHS, SinglecellHaystack; LR, Logistic regression; NegBinom, Negative binominal; Wilcoxon, Wilcoxon rank− sum test. **b** Line plot showing the runtimes of different methods applied to datasets with different sample size.

Finally, we compared the running time for the 10 methods. Monocle has the longest runtimes, for example, runtime on 5000 cells for Monocle is about 1 hour (Fig. 4b). In contrast, DEAPLOG, Biomod and t.test take below 1 min on 5000 cells. Moreover, the runtimes of Monocle, Negbinom, Poisson and MAST increase nonlinearly with the increase of cell number, rendering these tools powerless against big scRNA data analysis (Fig. 4b). We tested DEAPLOG on 57,000 cells, and the runtimes is only about 8 minutes. This further proves that DEAPLOG can be applied to big scRNA data analysis.

To sum up, DEAPLOG has high performance in robustness and accuracy at varying cell number, cell clusters and batches compared to other available tools. The short running times further makes it a reliable and attractive approach to identifying DEGs in even large-scale scRNA data.

### DEAPLOG has good performance in identifying DEGs on real scRNA-seq data

To evaluate the performance of our approach on real data, we applied DEAPLOG on a data from peripheral blood mononuclear cells (PBMC) based on 10X sequencing technologies, which is available on the 10X platform (see Method section). This data contains 6,280 cells and 19,869 genes after preprocessing and cleaning, and the UMAP plotting shows 11 cell clusters (Fig. 5a top-left panel). We use DEAPLOG to identify DEGs for each cell cluster in order to annotate them. The results show that we are able to identify different number of DEGs for each cluster although some of them (i.e. C8, C9 and C10) contain small numbers of cells (Fig. 5a). And indeed, we can annotate each cell cluster solely based on the top 10 DEGs identified by DEAPLOG as most of them are well-known markers for various immune cells (CD8A for CD8^+^ T cell, S100A8 for CD14^+^ monocytes (CD14^+^ Mono), CD79B for B cell, TRGV9 for natural killer T cell (NKT), GNLY for natural killer cell (NK), FCGR3A for CD16+ monocytes (CD16^+^ Mono), PPBP for megakaryocyte (Mega), PLD4 for plasmacytoid dendritic cell (pDC) and FCER1A for monocyte derived dendritic cell (mDC), see Supplementary Fig. 5A). Functional enrichment analysis of DEGs for each cell cluster further proves that DEGs identified by DEAPLOG are biological significant (see Supplementary Fig. 5B). Intriguingly, cluster C1 and C2, which we annotate as Naïve CD4^+^ (Naïve CD4^+^T) and CD8^+^T cells (Naïve CD8^+^T)^22^, also highly express many ribosomal genes based on their DEGs (Fig. 5a). This is especially true in Naïve CD4^+^T cells. To rule out the potential false-positive causes, we check the expression of these ribosomal genes in UMAP plot and confirm their expression specificity, suggesting that expressing of ribosomal genes is an intrinsic property of naïve T cells (Supplementary Fig. 5C). Indeed, this is very-well consistent with previous report that naïve T cells maintain a surprisingly large number of idling ribosomes to mount a rapid immune response, and provide a resource for protein turnover^23^. Overall, these results demonstrated that DEAPLOG has good performance in identifying DEGs on real scRNA-seq data.

**Fig. 5.**
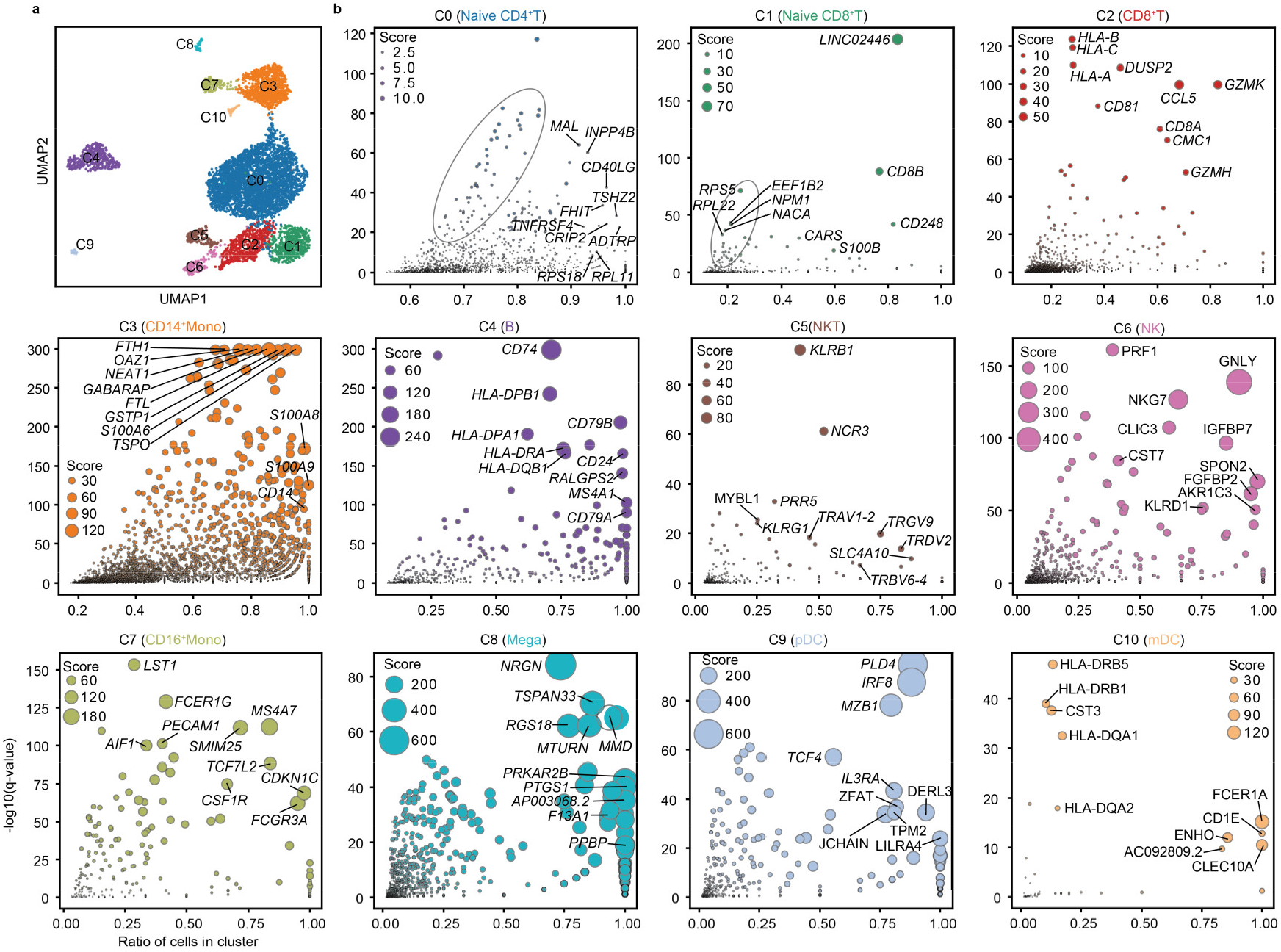
DEG identification by DEAPLOG applied on scRNA-seq data of peripheral blood mononuclear cells (PBMC). **a** UMAP plot showing the 11 cell clusters of PBMC data. **b** Scatter plots showing the p-value and ratio of all genes in each cell clusters, separately. Ratio represents the percentage of the cell signatures set for a gene in the given cell cluster.

### DEAPLOG is a novel tool can be applied in pseudo-temporal locating and ordering TFs that regulate cell fate decision

To evaluate the performance of DEAPLOG for pseud-temporal locating and ordering of genes, we applied DEAPLOG on the scRNA data from HSPC used in previous sections to explore the key TFs that regulate all lineages development. To infer the pseudo-times and locate the coordinates of genes, the pseudotime and embedding coordinates of cells are needed. We use two dimensions of embedding coordinates from UMAP as the coordinates of cells, and use pseudo-time calculated by ‘dpt’ function in Scanpy as the pseudo-time of cells. First, we identify DEGs for all cell clusters by DEAPLOG with ratio above 0.2 and adjusted p-value below 0.01 to narrow down the scope of gene list. We then extract all TFs from DEGs to infer the pseudo-times and spatial location of these TFs in UMAP^24^.

In total 1,241 TFs were identified in HSC lineages and about 71.6% of TFs are located at region of HSC/MPP (Fig. 6) suggesting that stemness, function, and fate of stem cells are exquisitely regulated by large number of TFs. Based on the pseudo-time and spatial location of TFs, we can also clearly identify those TFs involved in lymphoid, myeloid, and erythroid lineages development in a sequentially activated manner from HSC to differentiated cells. For example, *MECOM, HOPX, HLF* and *MLLT3* located in top, middle, left bottom and right bottom region of HSC/MPP population, respectively, which suggests that *MECOM* and *HOPX* are more likely function in maintaining the stemness of HSC/MPP, *HLF* and *MLLT3* are in driving the development of lymphoid and erythroid lineage development. Indeed, Lin et.al reported that HOPX is crucial in maintaining quiescent of HSC through CXCL12 pathway in mice^25^. Besides, Tang et.al reported that *Hlf* expression marks early emergence of HSC precursors in mice and Pina et.al reported that MLLT3 regulates early human erythroid and megakaryocytic cell fate^26,27^. All supports the above inference. More importantly, through DEAPLOG, we can construct pseudo-time chains of TFs that regulate cell differentiation from stem cell into terminally differentiated cell. For example, the pseudo-time chain of MLLT3-PBX1-TRIB2 likely regulates the differentiation process of megakaryocyte and erythrocyte from HSC/MPP, and the pseudo-time chain of HLF-GAS7-RAG1 is more likely to regulate the differentiation process of B cell from HSC/MPP. Therefore, we demonstrate with real data that DEAPLOG can accurately infer the pseudo-time and locate embedding coordinate of genes and construct pseudo-time chains of TFs that drive development of lineages.

**Fig. 6.**
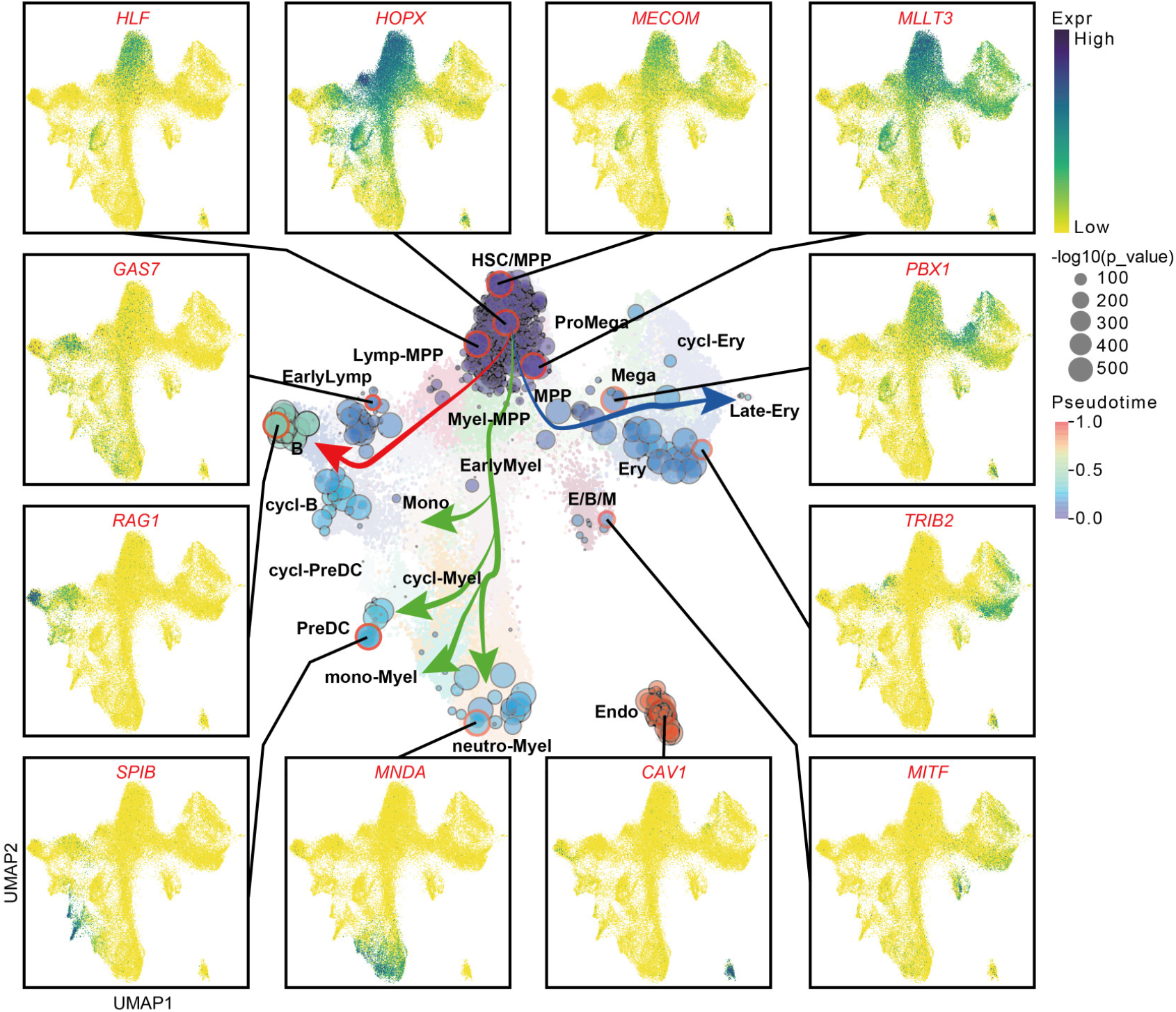
Pseudo-temporal locating and ordering of transcript factors (TFs) by DEAPLOG applied on scRNA-seq data of hematopoietic stem/progenitors (HSPC) and its derived cells. UMAP plot showing the pseudo-location and -time of all TFs enriched by DEAPLOG and the normalized and logarithmic expression level of the 12 representative TFs. HSC, hematopoietic stem cells; MPP, multipotent progenitor; Lymp-MPP, lymphoid MPP; Myel-MPP, myeloid MPP; Mono, monocyte; cycl-Myel, cycling lymphoid cell; PreDC, dendritic progenitor; Neutro-Myel, neutrophil-myeloid cell; Mega, megakaryocytic cell; PreMega, megakaryocytic progenitor; Ery, erythroid cell; Late-Ery, lately Ery; E/B/M, eosinophil/basophils/mast cells; Endo, endothelial cell.

### DEAPLOG can be used in identifying DEGs in spatial transcriptomic data

To explore whether DEAPLOG can be used to identify DEGs in spatial transcriptomic data, we used the slice dataset of mouse brain sagittal posterior, which is available from 10x Visium platform (see Method section). This dataset contains gene expression information of 3353 spots, 2D spatial coordinates of the sports within the tissue and the original tissue slice image. We use stlearn package to preprocess, reduce dimension, cluster and visualize the transcriptomic data^28^. Finally, the whole spots are clustered into 18 clusters (Fig. 7a and b).

**Fig. 7.**
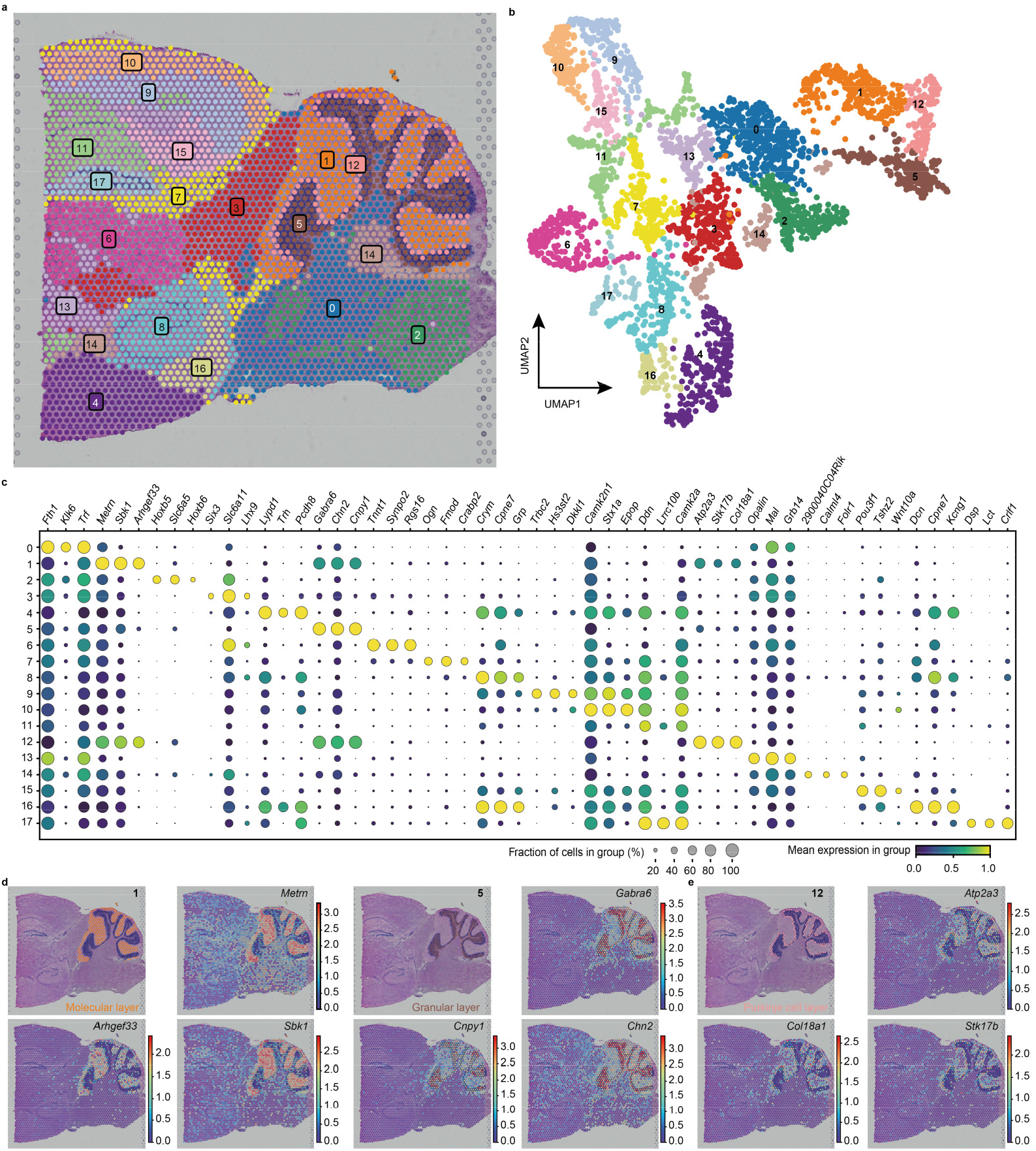
DEG identification by DEAPLOG applied on spatial transcriptomic data of mouse brain sagittal posterior. **a** Spatially resolved heatmaps across tissue sections showing the spatial distribution of spots clusters. **b** UMAP plot showing the spot clusters. **c** Dot plot showing the top three DEGs identified by DEAPLOG for each spot cluster. **d** Spatially resolved heatmaps across tissue sections showing the normalized expression pattern of the top three DEGs expressed in molecular layer (left panel, cluster 1), granular layer (middle panel, cluster 5) and Purkinje cell layer (right panel, cluster 12) of cerebellar cortex.

DEAPLOG successfully identify DEGs for all 18 clusters and the top three DEGs of each cluster are very specific (Fig. 7c). To verify that these DEGs are also specific in spatial coordinates, we selected one gene for each spot cluster from DEGs and mapped these gene into spatial coordinates. The results show that these genes had specific spatial expression patterns even in very fine spatial regions (see Supplementary Fig 6A). For example, *Metrn, Sbk1 and Arhgef33* are specifically expressed in the molecular layer of cerebellar cortex, meanwhile, *Gabra6, Chn2 and Cnpy1* are specifically expressed in the granular layers of cerebellar cortex in spatial coordinates^29,30^ (Fig. 7d). In addition, DEAPLOG identified three genes, *Atp2a3, Stk17b*, and *Col18a1*, specifically expressed in a narrow region, which locates between the molecular layer and granular layer of the cerebellar cortex (Fig. 7e). It has been reported that the three genes regulate the function of Purkinje cells^31-33^. Therefore, the specific expression patterns of these three genes indicate that this narrow region (cluster 12) is composed of Purkinje cells, which is consistent with the existing reports that the mature cerebellar cortex is composed of three layers: inner granular layer, Purkinje cell layer and molecular layer^34^. The results of in situ hybridization (ISH) further confirmed that Atp2a3, Stk17b, and Col18a1 can be the marker genes of Purkinje cells^35^ (see Supplementary Fig 6B). In conclusion, DEAPLOG can be also used to identify spatially DEGs on spatial transcriptomic data.

## Discussion

DEGs identification is a key step of scRNA-seq data analysis, however, the available tool for it is limited. We present a new method, DEAPLOG, by combining the algorithms of polynomial fitting and hypergeometric distribution test, rather than probability distribution that the current methods generally use. With both simulated and real scRNA-seq data, we demonstrate that our method has superior performance than existing tools in one or multiple aspects of accuracy, robustness and efficiency. While DEAPLOG excels at differential expression analysis of scRNA-seq data between cell categories, it can be easily extended to other variables, such as tissue types, disease status, time series, and even spatial locations. Notably, we recommend that DEGs should be identified for each cell clusters when performing other condition of tests, considering the heterogeneity of gene expression in cell clusters. As our method does not based on the probability distribution, it eliminates the obsession of the “droupout event” and the multimodal distribution of genes in scRNA-seq data. Considering recent reports that DESeq2 and edgeR have unexpectedly high false discovery rates when processing population-level RNA-seq studies with large sample sizes, we suggest that DEAPLOG can be a replacement for these two methods or even the Wilcoxon rank-sum test which was recommended by authors to perform differential expression analysis for population-level RNA-seq studies with large sample sizes^36^. Because, like the Wilcoxon rank-sum test, DEAPLOG is also more concerned with the sequencing of gene expression than with the actual expression value.

Another merit of DEAPLOG is can be used to infer the pseudo-time and location of genes, hence to construct pseudo-time chains of regulators that drive development of lineages, which has important implications in all aspects of cell fate decision analysis. Finally, DEAPLOG uses anndata as input data format, which can work seamlessly with popular single-cell analysis platforms, such as Scanpy. All above makes DEAPLOG a reliable and attractive approach in scRNA data analysis. Indeed, we have successfully applied it in large-scale human embryonic limb development data analysis and several other applications^37-39^.

Despite using a completely different framework to identify DEGs, DEAPLOG still returns inflated p-values like many existing methods. Therefore, DEAPLOG identifies a large number of statistically significant DEGs, which lead to a certain percentage of FPR. In order to reduce the FPR, on one hand, we use the Benjamini-Hochberg method to correct the p-values. on the other hand, we added a parameter “ratio” to characterize the expression ratio of a certain gene in given cell cluster. We recommend setting the threshold of “ratio” to 0.2 and adjusted p-values to 1e-30 to reduce the number of DEGs. Additionally, the threshold of “ratio” can be set by the user, allowing to identify genes that have multiple clusters distribution. To identify the cell signature for a gene, we define the point with the maximum curvature on the fitting curve of a gene expression as threshold. With it, the cells before this point are defined as the cell signature of this gene. However, we realized that in extreme cases, the fitting curve around the threshold point might have low slope (slowly curve descending), which makes it naturally unreasonable to defined cells before this point as the cell signature of a gene. This may be another reason leads to FPR.

## Methods

### DEAPLOG methodology

DEAPLOG consists of three functions: (i) get_DEG_uniq to identify genes that are differentially expressed in only one cell type; (ii) get_DEG_multi to identify genes that are differentially expressed in one or more cell types; (iii) get_genes_location_pseudotime to infer the pseudo-time and embedding coordinates of genes based on pseudo-time and embedding coordinates of cells.

For the first two functions, DEAPLOG combines polynomial fitting and hypergeometric distribution test to identify DEGs. To determine whether a gene G is DEG, first, we rank all cells in descending order based on the expression levels of gene G and use the ranked numbers of the cells (from 1 to N, N is total number of cells) as the independent variables and the expression values of gene G as the dependent variables to characterize the expression pattern of gene G. Then we use polynomials to fit the expression pattern of gene G. The fitting function is defined as follow:

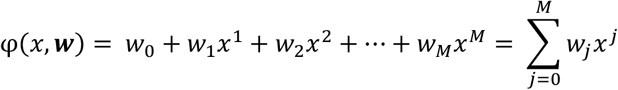

where *x* represents ranked number of the cell, *w*_*j*_ represents the coefficient of *x*^*j*^ and *M* is the highest degree of the polynomial. According to the least square principle, this loss function is defined as follow:

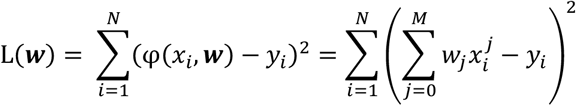

Where N is the total number of cells and *y*_*i*_ is the expression value of the gene in cell *i*. The least square method is to minimize L(***w***), that is, the partial derivative of L(***w***) to each *w* is 0:

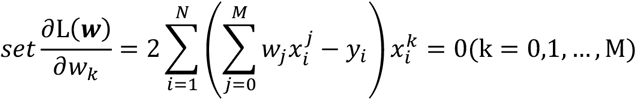

Then, solving the above equation to get each coefficient ***w***. To determine the best value of *M*, we used the standard deviation of the fitting residual to measure the accuracy of the polynomial fitting. For *N* cells and the degree *M*, the degree of freedom of the fitting residual data is N-M-1, therefore, the standard deviation of the fitting residual is defined as follow:

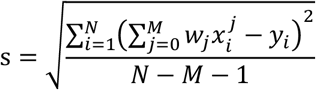

The degree corresponding to the minimum value of s is defined as the best value of *M*. When the fitted curve is determined, we used the curvature formula to determine the *x*_*k*_ corresponding to the maximum curvature of the curve, and selected the top *x*_*k*_ cells as the cell signature of gene G. The curvature formula is as follow:

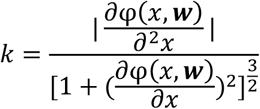

Thus, let the first derivative of *k* be 0 to get the *x*_*k*_:

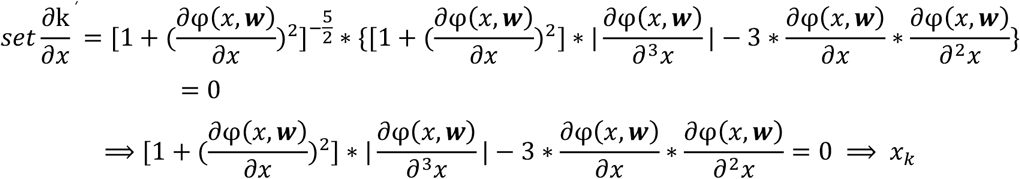

For each gene, we performed the above process to get the cell signature for each gene and constructed a cell signature sets for all genes. After that, we applied the hypergeometric distribution test to determine which cell clusters these cell signature sets belong to and further to determine whether the gene corresponding to cell signature set is the DEG of the cell cluster. the probability that a cell signature set belongs to a cell cluster was defined as follow:

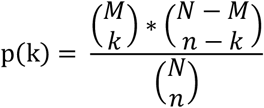

The *p*-value of hypergeometric test was defined as follow:

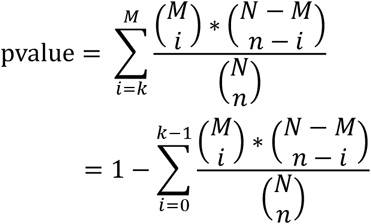

Where *N* represents the number of all cells, *M* represents the cell number of the cell cluster, *n* represents the cell number of the cell signature set, *k* represents the number of cells that belong to both the cell cluster and the cell signature set. Then, the Benjamini-Hochberg (BH) method was used to correct *p*-values. Additionally, we defined two variables: ‘ratio’ and ‘score’ as follows:

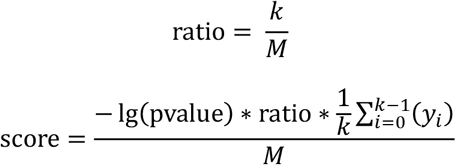

The ‘ratio’ reflects the proportion of cells that belong to both the cell cluster and the cell signature set. *y*_*i*_ is the expression value of the gene in cell *i* which belong to intersection of the cell cluster and the cell signature set. The ‘score’ reflects the specificity of a gene G as the DEG in cell cluster. For inferring the pseudo-time and embedding coordinates of genes, we defined the pseudotime and embedding coordinate of a gene as the follows:

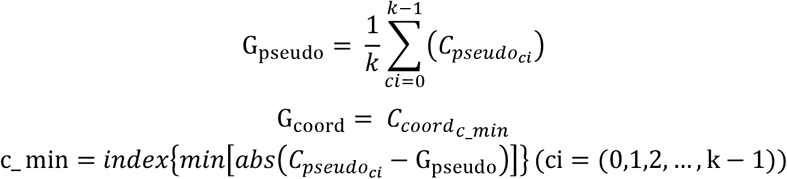

Where *k* represents the number of cells that belong to both the cell cluster and the cell signature set, 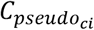 represents the pseudotime of cell *ci* which belongs to both the cell cluster and the cell signature set.

### Comparison of methods on artificial datasets

We use Splatter (v 1.14.1) to simulate 27 artificial datasets of varying cell number, cell clusters and batches using Splatter. 6 datasets contain 50, 100, 500, 1000, 2500 and 5000 cells, separately and each dataset consists of 2 cell clusters; 5 datasets contain 2, 5, 10, 25 and 50 cell clusters with fixed 200 cells per cluster, separately; 5 datasets contain 2, 5, 10, 25 and 50 cell clusters separately and each dataset contains 10,000 cells; 5 datasets of which the cell number in cell clusters follows Poisson, Beta, Gamma, Normal and Uniform distribution, respectively and each dataset contains 10k cells with 10 clusters; 6 datasets introduced 1, 2, 4, 6, 8, 10 batch effects, separately and each dataset contains 5000 cells in 10 clusters. The output of Splatter contains differential expression factors, showing whether a gene has differential expression (factor different from 1) or not (factor = 1) in each cell cluster. For the 6 artificial datasets of varying cell number, the proportion of DGEs was set at 10% for each cell cluster. For the rest 21 datasets, the proportion of DGEs was set at 5% for each cell cluster.

We apply our approach, SCHS (v 0.3.4), monocle (v 2.20.0), as well as approaches (bimod, logreg, MAST, negbinom, poisson, t.test, wilcox) available through the FindAllMarkers function of the Seurat (4.1.1) toolkit on these artificial datasets. For 9 existing methods we predict DEGs by comparing each cluster versus all other clusters with default options and without the filtering. All evaluated methods assign p-values to genes reflecting their degree of differential expression. Finally, for each DEG prediction method, we calculate the AUC under the ROC curve for every artificial dataset using the pROC (v 1.17.0.1) package in R.

### Data filtering and preprocessing of real scRNA-seq datasets

The data filtering and preprocessing of real scRNA-seq datasets is performed by using the Python package, Scanpy (v.1.9.1). We download the processed data of the scRNA-seq data of human cells derived from HSPCs, which contains the raw counts matrix, the information of cell clusters and the embedding coordinates of TSNE and UMAP. To identify the DEGs by DEAPLOG, we normalize and logarithmic transform the raw counts matrix. To calculate the pseudo-time of cells, first, we select 1,984 highly variable genes (HVGs) by using the “pp.highly_variable_genes” function in Scanpy with the parameter: batch_key = “sampleID”. Secondly, principal component analysis (PCA) is conducted using the 1,984 HVGs and the first 42 PCs are used for downstream analysis. Finally, the pseudo-time of cells is calculated by using the ‘diffmap’ and ‘dpt’ functions in Scanpy with default parameters.

For the scRNA-seq data of PBMCs, we download the raw count matrix without any filtering. Filtering retained cells with < 20k total unique molecular identifiers (UMIs), gene number between 400 and 4000, <8% UMIs mapped to mitochondrial genes, and retained genes expressed in at least 3 cells. We also filtered cells with <0.1 BH-pvalue by using scrublet package [PMID: 30954476]. After that, we normalize and logarithmic transform the raw counts matrix and select 2,447 HVGs by using the “pp.highly_variable_genes” function with the default parameter. Then, we regress out the cell cycle, total counts and expression of mitochondrial genes by using the“pp.regress_out “ function and compute the neighborhood graph by using the “pp.neipgbors” function with 41 PCs. The further dimensionality reduction and clustering are performed by using the “tl.umap” and the “tl.leiden” with default parameters, separately.

### Data filtering, preprocessing and visualization of real spatial transcriptomic data

The filtering, preprocessing and visualization of spatial transcriptomic data for mouse brain sagittal posterior is performed by using the Python package, stlearn (v.0.3.2). For spot image, we firstly tile hematoxylin-eosin staining (H&E) image to small tiles based on spatial location by using “pp.tiling” function and then extract latent morphological features from the small tiles by using “pp.extract_feature” function with default parameters. For gene matrix, we firstly normalize the data based on the spatial location, tissue morphological feature that has been extracted from the small tiles and gene expression information by using “spatial.SME.SME_normalize” function with the raw counts and the parameter, weight=“weights_matrix_pd_md” after filtering genes that didn’t expressed in any spot and then scale the data by using the “pp.scale” function. After that, we perform preliminary dimension reduction and compute a neighborhood graph by using the “em.run_pca” and the “pp.neighbor” functions with the 50 PCs and the parameter, n_neighbor=20, separately. The further dimensionality reduction and clustering are performed by using the “tl.umap” and the “tl.clustering.louvain” with default parameters, separately.

## Data availability

The 27 artificial single-cell datasets are generated by using Splatter are available in figshare (https://figshare.com/account/articles/21383190). The scRNA-seq data of HSPCs is downloaded from GEO (https://www.ncbi.nlm.nih.gov/geo/query/acc.cgi?acc=GSE155259); The scRNA-seq data of 10k PBMC and the spatial transcriptomic data of mouse brain are downloaded from 10x Genomics (https://www.10xgenomics.com/resources/datasets/10-k-human-pbm-cs-5-v-2-0-chromium-controller-2-standard-6-1-0; https://www.10xgenomics.com/resources/datasets/mouse-brain-serial-section-2-sagittal-posterior-1-standard)

## Code availability

DEAPLOG is implemented as a Python package, available from PyPI (https://pypi.org/project/deaplog) and GitHub (https://github.com/ZhangHongbo-Lab/DEAPLOG). The repository includes additional instructions for installation in Python and example applications. Additionally, the notebooks for simulating artificial datasets by Splatter, running 10 existing methods to predict DEGs, preprocessing real scRNA-seq datasets and spatial dataset are available from GitHub (https://github.com/ZhangHongbo-Lab/DEAPLOG/notebook/).

## Acknowledgements

We thank the members of the laboratories of Prof. Hongbo Zhang (Sun Yat-sen University), Prof. Miaoxin Li (Sun Yat-sen University) and Prof. Sarah A Teichmann (Wellcome Sanger Institute) for helpful discussions. This work was supported by the National Key R&D Program (grant number: 2022YFA1104904 and 2019YFA0801703) and the National Natural Science Foundation of China (grant numbers: 31871370).

## Author contributions

B.Z. conceived of the project and methodology, implemented the methods and performed the analyses. H.Z. supervised the project. B.Z. and H. Z. wrote the manuscript and approved the submitted version.

