## Supplemental Figures for "A method for differential expression analysis and pseudo-temporal locating and ordering of genes in single-cell transcriptomic data"

Supplementary Figures:

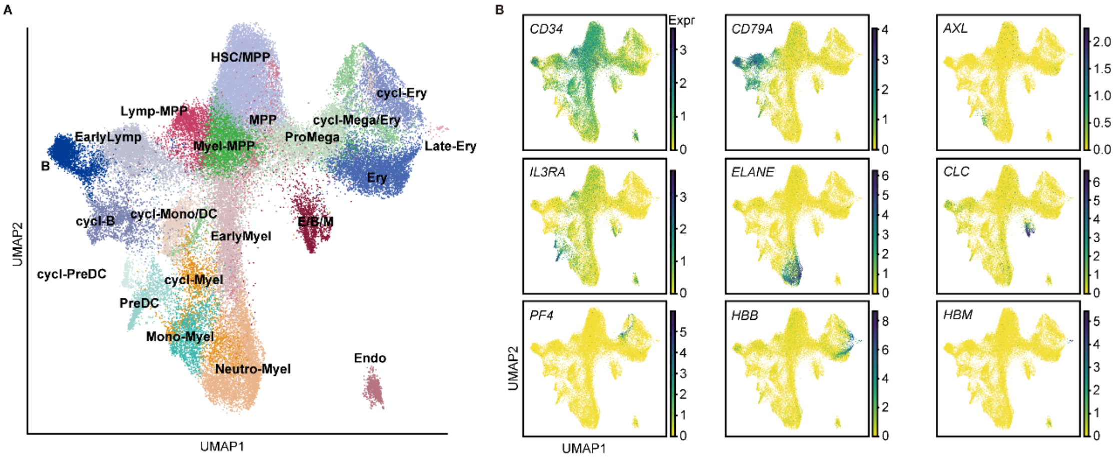


**Supplementary Fig. 1 Cell atlas of scRNA-seq data of hematopoietic stem/progenitors (HSPC) and its derived cells**. **A** Umap plot showing the cell clusters of cell atlas. HSC, hematopoietic stem cells; MPP, multipotent progenitor; Lymp-MPP, lymphoid MPP; Myel-MPP, myeloid MPP ; Mono, monocyte; cycl-Myel, cycling lymphoid cell; PreDC, dendritic progenitor; Neutro-Myel, neutrophil-myeloid cell; Mega, megakaryocytic cell; PreMega, megakaryocytic progenitor; Ery, erythroid cell; Late-Ery, lately Ery; E/B/M, eosinophil/basophils/mast cells; Endo, endothelial cell. **B** Umap plot showing the normalized and logarithmic expression level of well-known marker genes for each cell cluster.

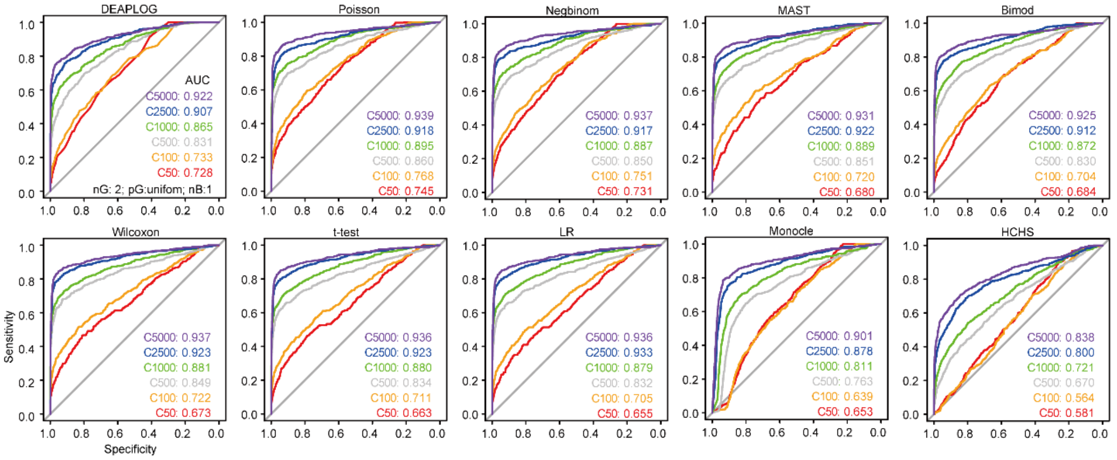


**Supplementary Fig. 2 Receiver operating characteristic (ROC) curves showing ability of DEAPLOG and other 9 methods to detect differentially expressed genes (DEGs) applied on the simulated 6 artificial datasets with different cell numbers.** Each dataset contains 2 cell clusters and the cell number in each cell cluster is same. AUC, area under curve; SCHS, SinglecellHaystack; LR, Logistic regression; NegBinom, Negative binominal; Wilcoxon, Wilcoxon rank−sum test.

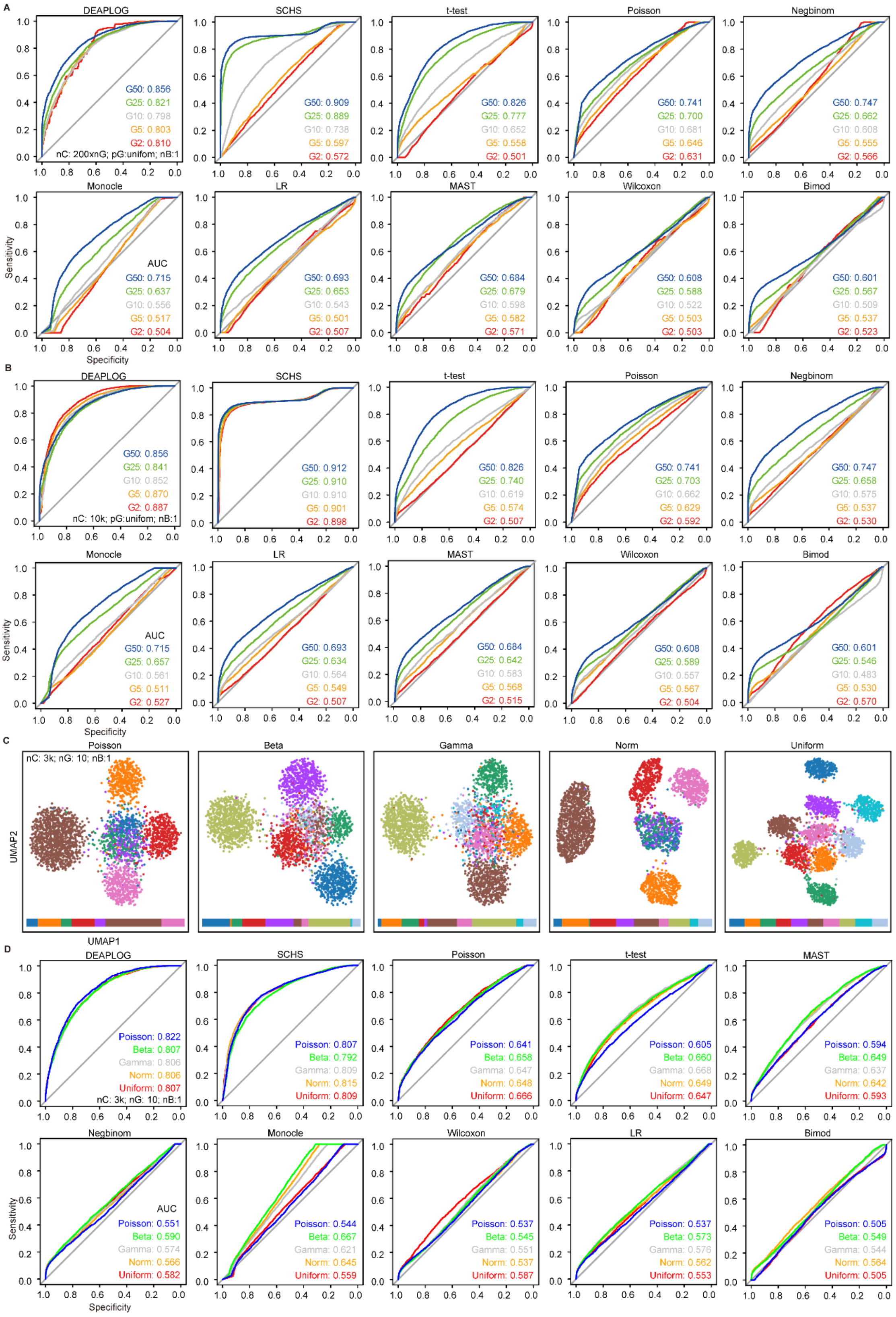


**Supplementary Fig. 3 Receiver operating characteristic (ROC) curves showing ability of DEAPLOG and other 9 methods to detect differentially expressed genes (DEGs) applied on the simulated 15 artificial datasets with different cell clusters. A** ROC curves showing ability of DEAPLOG and other 9 methods to detect DEGs applied on 5 datasets which contains 2, 5, 10, 25 and 50 cell clusters with fixed 200 cells per cluster in each dataset, separately. **B** ROC curves showing ability of DEAPLOG and other 9 methods to detect DEGs applied on 5 datasets which contains 2, 5, 10, 25 and 50 cell clusters with 10,000 total cell number per dataset, separately. **C** UMAP plots showing the cell cluster distribution for 5 datasets in which the number of cell clusters followed Poisson, Beta, Gamma, Normal and Uniform distribution, separately. **D** ROC curves showing ability of DEAPLOG and other 9 methods to detect DEGs applied on 5 datasets in **C**. AUC, area under curve; SCHS, SinglecellHaystack; LR, Logistic regression; NegBinom, Negative binominal; Wilcoxon, Wilcoxon rank−sum test.

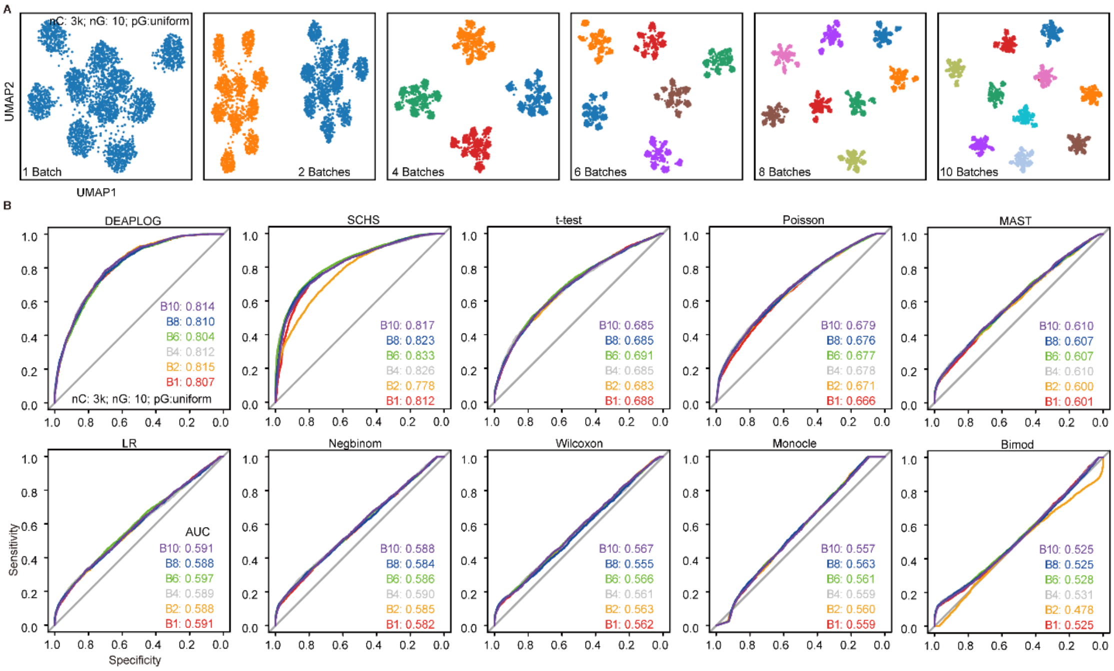


**Supplementary Fig. 4 Receiver operating characteristic (ROC) curves showing ability of DEAPLOG and other 9 methods to detect differentially expressed genes (DEGs) applied on the simulated 6 artificial datasets with different batches.** **A** UMAP plots showing the batches for 5 datasets with 1, 2, 4, 6, 8 and 10 batches, separately. **B** ROC curves showing ability of DEAPLOG and other 9 methods to detect DEGs applied on 5 datasets in **A**. AUC, area under curve; SCHS, SinglecellHaystack; LR, Logistic regression; NegBinom, Negative binominal; Wilcoxon, Wilcoxon rank−sum test.

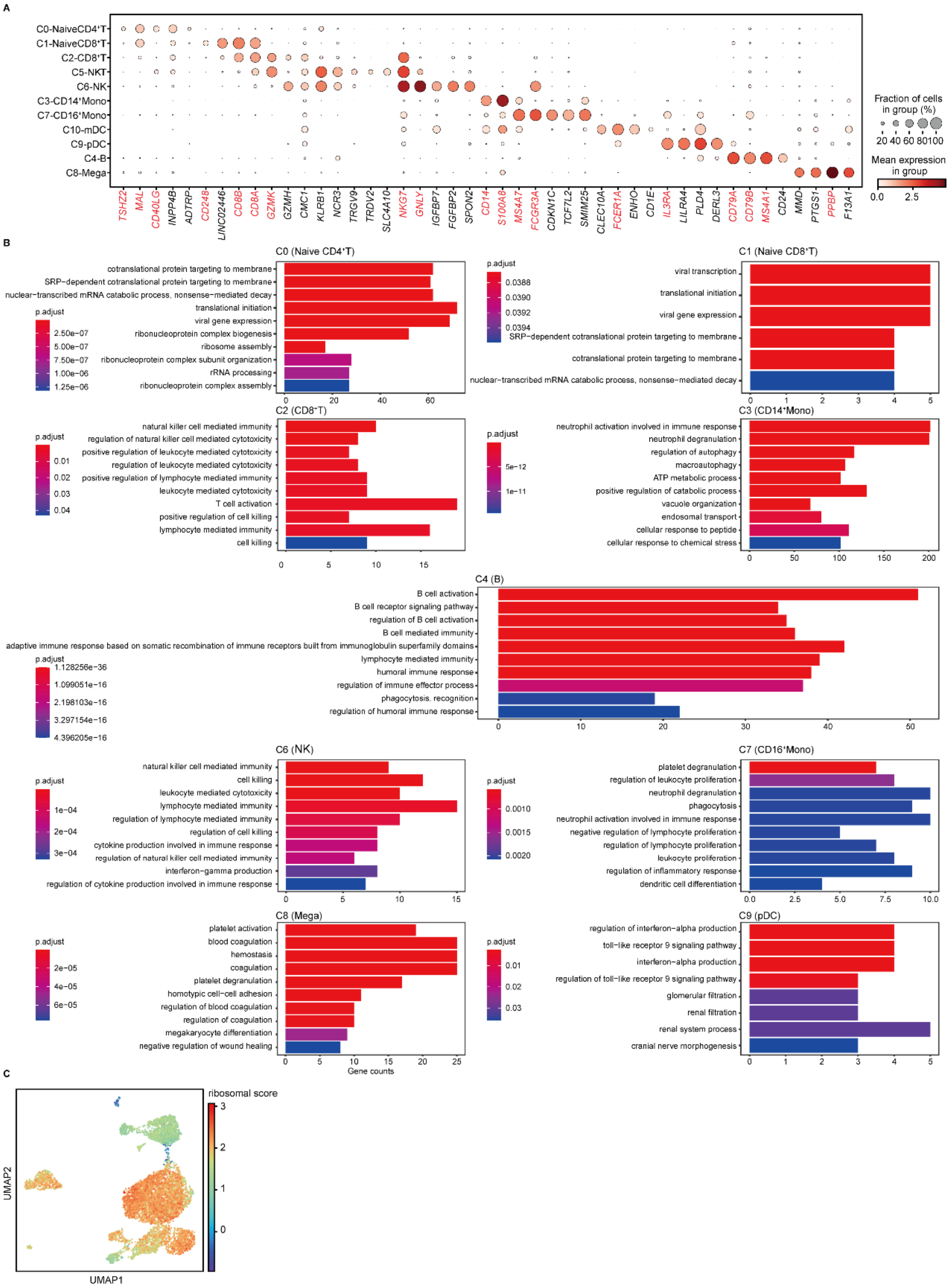

**Supplementary Fig. 5 DEGs identified by DEAPLOG and its Gene Ontology (GO) enrichment analysis of scRNA-seq data from peripheral blood mononuclear cells (PBMC). A** Dot plots showing the normalized and logarithmic expression level of top DEGs identified by DEAPLOG for each cell cluster. Genes colored by red are well-known marker genes for each cell cluster. **B** bar charts showing the significant biological process (BP) terms of GO for DEGs identified by DEAPLOG for each cell cluster. **C** UMAP plot showing the expression score of ribosomal genes.

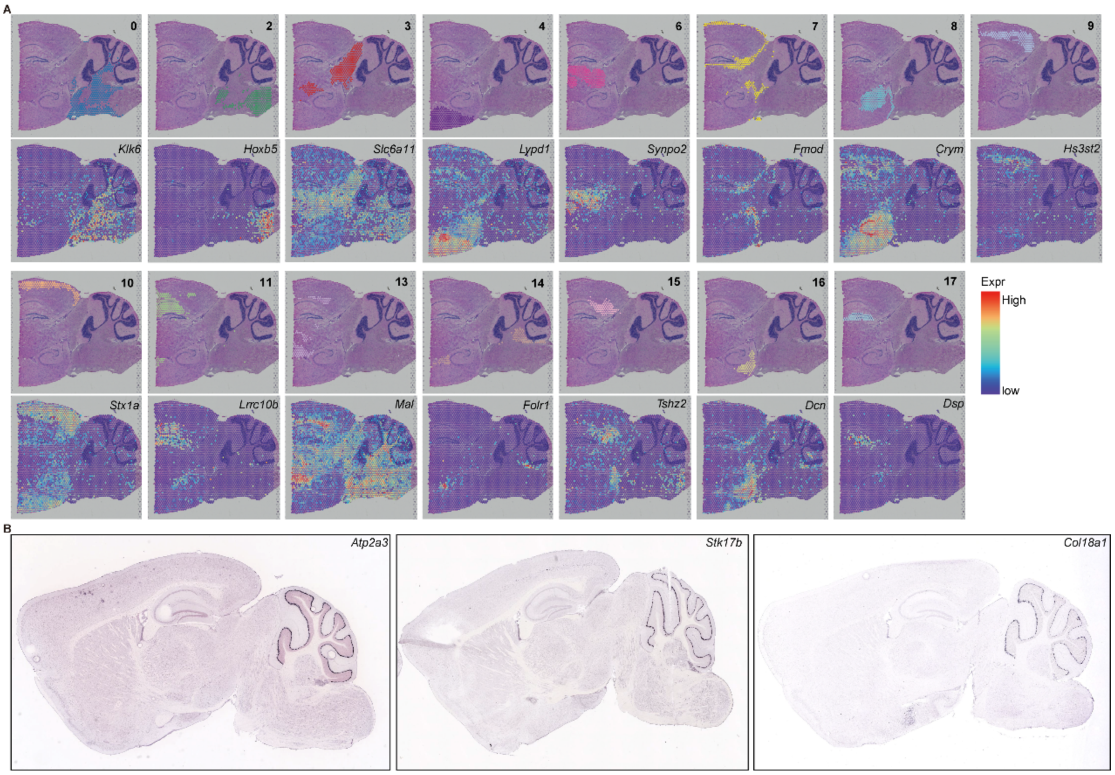


**Supplementary Fig. 6 The expression pattern of selected DEGs in spatial transcriptome of mouse brain sagittal posterior.** **A** Spatially resolved heatmaps across tissue sections showing the normalized expression pattern of selected DEGs for each spot cluster. **B** Expression of *Atp2a3* (https://mouse.brain-map.org/gene/show/32793), *Stk17b* (https://mouse.brain-map.org/gene/show/62437), and *Col18a1*(https://mouse.brain-map.org/gene/show/12605) in mouse brain sagittal posterior.
